# ESCRT recruitment by the inner nuclear membrane protein Heh1 is regulated by Hub1-mediated alternative splicing

**DOI:** 10.1101/2020.06.25.171694

**Authors:** Matías Capella, Lucía Martín Caballero, Boris Pfander, Sigurd Braun, Stefan Jentsch

**Affiliations:** Molecular Cell Biology, Max Planck Institute of Biochemistry, Martinsried, Germany; Department of Physiological Chemistry, Biomedical Center (BMC), Ludwig Maximilians University of Munich, Martinsried, Germany; International Max Planck Research School for Molecular and Cellular Life Sciences, Martinsried, Germany; DNA Replication and Genome Integrity, Max Planck Institute of Biochemistry, Martinsried, Germany

**Keywords:** ESCRT, LEM domain, nuclear envelope surveillance, nuclear pore complex, splicing

## Abstract

Misassembled nuclear pore complexes (NPCs) are removed by sealing off the surrounding nuclear envelope (NE), which is mediated by members of the ESCRT (<u>e</u>ndosomal <u>s</u>orting <u>c</u>omplexes <u>r</u>equired for <u>t</u>ransport) machinery. Recruitment of ESCRT proteins to the NE is mediated by the interaction between the ESCRT member Chm7 and the inner nuclear membrane protein Heh1, which belongs to the conserved LEM family. Increased ESCRT recruitment results in excessive membrane scission at damage sites but its regulation remains poorly understood. Here, we show that Hub1-mediated alternative splicing of *HEH1* pre-mRNA, resulting into its shorter form Heh1-S, is critical for the integrity of the NE. ESCRT-III mutants lacking Hub1 or Heh1-S display severe growth defects and accumulate improperly assembled NPCs. This depends on the interaction of Chm7 with the conserved MSC domain only present in the longer spliced variant Heh1-L. Heh1 variants assemble into heterodimers and we demonstrate that a unique splice segment in Heh1-S suppresses growth defects associated with uncontrolled interaction between Heh1-L and Chm7. Together, our findings reveal that Hub1-mediated splicing generates Heh1-S to regulate ESCRT recruitment to the nuclear envelope.

**Summary statement:** Heh1-S, the Hub1-mediated spliced version of *HEH1* pre-mRNA, contributes to nuclear envelope maintenance by preventing excessive recruitment of Chm7.

## Introduction

In eukaryotic cells, the double lipid bilayer membrane forming the nuclear envelope (NE) physically separates the nucleoplasm from the cytoplasm. Translocation of macromolecules in and out of the nucleus is mediated by nuclear pore complexes (NPCs), which are composed of multiple copies of ∼30 different protein components called nucleoporins (Alber et al., 2007; Cronshaw et al., 2002; D’Angelo et al., 2008; Rout et al., 2000; Strambio-De-Castillia et al., 2010). Notably, disruption of the precise structure of NPCs has been linked to various diseases, including cancer (Nofrini et al., 2016; Sakuma and D’Angelo, 2017).

The <u>e</u>ndosomal <u>s</u>orting <u>c</u>omplexes <u>r</u>equired for <u>t</u>ransport (ESCRT) components have been previously shown to be involved in the quality control of NPC assembly (Webster et al., 2014, 2016). The ESCRT machinery comprises five evolutionarily conserved complexes, ESCRT-0, I, II, III and the AAA ATPase Vps4. Together, they orchestrate membrane remodeling and scission in diverse cellular processes including receptor sorting, virus budding, cytokinesis and plasma membrane wound repair (Alonso Y Adell et al., 2016; Hurley, 2015; Vietri et al., 2020). In addition, the ESCRT machinery has been implicated in nuclear functions. For instance, the ESCRT machinery coordinates resealing of nuclear envelope fragments in humans and fission yeast after cell division (Lee et al., 2020; Olmos et al., 2015; Pieper et al., 2020; Vietri et al., 2015). Moreover, yeast mutants defective in either ESCRT-III or Vps4 accumulate misassembled NPCs at specific NE areas coined SINC (<u>s</u>torage of <u>i</u>mproperly assembled <u>n</u>uclear pore <u>c</u>omplexes compartment; Webster et al., 2014). While ESCRT-dependent surveillance is crucial for maintaining nuclear compartmentalization, uncontrolled recruitment to the NE can be detrimental for cell viability (Gu et al., 2017; Thaller et al., 2019; Willan et al., 2019), suggesting that this process needs to be tightly regulated. However, how excessive ESCRT recruitment to NE is prevented remains unknown.

The ESCRT machinery is recruited to the NE through interaction of the ESCRT-III adaptor CHMP7 (Chm7 in budding yeast) with the inner nuclear envelope protein LEM2 (Gu et al., 2017; Halfmann et al., 2019; Lee et al., 2020; Pieper et al., 2020; Thaller et al., 2019; von Appen et al., 2020; Webster et al., 2016). LEM2 and its paralog MAN1 belong to the conserved LEM (<u>L</u>ap2, <u>e</u>merin, <u>M</u>AN1)-domain protein family. They comprise a LEM and a winged-helix MSC domain (<u>M</u>AN1-<u>S</u>rc1 <u>C</u>-terminal) at their N- and C-terminus, respectively, which face the nucleoplasm and are separated by two transmembrane domains (TM; Brachner and Foisner, 2011). While the LEM domain has been the focus of many studies, several important functions have been recently reported for the MSC domain. For instance, LEM2 anchors telomeres and silences heterochromatin through its MSC domain; it further recruits CHMP7 to seal NE holes (Barrales et al., 2016; Gu et al., 2017; Halfmann et al., 2019; Hirano et al., 2018; Pieper et al., 2020; Thaller et al., 2019; von Appen et al., 2020). In budding yeast, the LEM2 homolog Heh1 (also known as Src1) interacts with Chm7 as well (Thaller et al., 2019; Webster et al., 2016). Interestingly, *HEH1* is a rare example of alternative splicing in *Saccharomyces cerevisiae*, resulting in two variants: a long form of 834 aa (Heh1-L) and a short form of 687 aa (Heh1-S; Fig S1A). Alternative splicing causes a shift in the open reading frame of Heh1-S, resulting in the appearance of 49 unique residues that replace the second TM and the C-terminal MSC domain (Grund et al., 2008; Rodríguez-Navarro et al., 2002; Fig. S1A). This splicing reaction is assisted by Hub1 (also known as UBL5), which belongs to the family of ubiquitin-like proteins but does not form conjugates (Karaduman et al., 2017; Lüders et al., 2003; Mishra et al., 2011; Wilkinson et al., 2004; Yashiroda and Tanaka, 2004). Cells lacking Hub1 express only Heh1-L (Karaduman et al., 2017; Mishra et al., 2011). While Heh1 has been described to contribute to NE integrity, DNA tethering and chromatin silencing (Chan et al., 2011; Grund et al., 2008; Mekhail et al., 2008; Webster et al., 2016), the physiological relevance of Hub1-mediated *HEH1* splicing remains largely unexplored. Here we examine the link between Hub1-mediated splicing and NE integrity. We demonstrate that Heh1-S is critical for normal cell growth when ESCRT-III subunits are absent. Lack of Heh1-S causes a severe growth phenotype due to a toxic gain-of-function of Heh1-L, resulting in increased interaction with Chm7. Moreover, we find that Heh1-S and Heh1-L form heterodimers and that the unique splice segment in Heh1-S is critical to suppress cellular toxicity. Together, we propose that Hub1-dependent alternative splicing modulates the binding of Chm7 to Heh1-L, thus avoiding excessive recruitment of Chm7 to the nuclear envelope.

## Results

### ESCRT-III deficiency is toxic in cells lacking Hub1

When overexpressed, Hub1 enhances splicing efficiency and activates several cryptic introns (Karaduman et al., 2017). In particular, Hub1 overexpression induces mis-splicing of Did4 (also known as Vps2), the major structural component of ESCRT-III together with Snf7 (Vps32; Babst et al., 2002; Karaduman et al., 2017). Mutations in Hub1 further exacerbate the growth defects associated with the deletion of the ESCRT-III subunit *VPS24* (Costanzo et al., 2016). Moreover, we identified several members of the ESCRT-III complex in a genetic screen for mutants displaying synthetic growth defects in combination with *hub1*Δ (our unpublished results). Together, these findings suggest a functional link between Hub1 and the ESCRT machinery. To further examine this relationship, we crossed a *hub1*Δ strain with a mutant lacking the ESCRT-III subunit Did4. Consistent with previous studies, we found that lack of Hub1 does not affect growth at 37°C. However, the additional deletion of *HUB1* enhanced the temperature sensitivity of the *did4*Δ mutant (Fig. 1A). Similarly, we observed a synthetic growth phenotype for cells lacking both Hub1 and Snf7. In contrast, no synthetic genetic interaction was seen for *hub1*Δ in combination with a deletion of the ESCRT-II component *VPS25* (Fig. 1B). These data suggest that Hub1 becomes essential when the ESCRT-III but not ESCRT-II complex is absent.

**Fig. 1.**
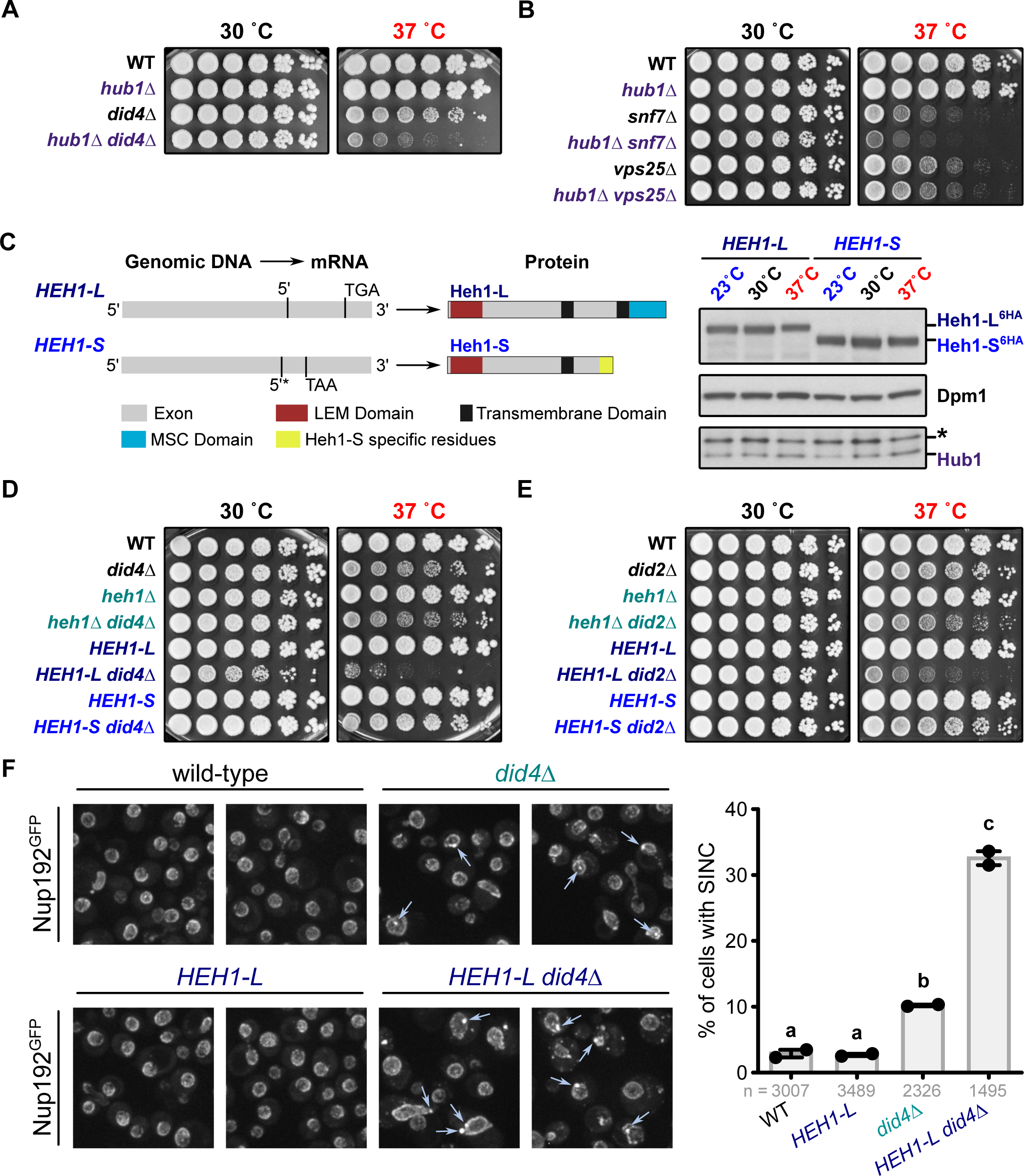
Cells lacking Heh1-S and ESCRT-III proteins display a synthetic growth phenotype and accumulate NPCs at the SINC. (A,B) Five-fold serial dilutions of the indicated strains were spotted and grown on YPD at 30°C and 37°C for 3 days. (C) Left panel: Scheme that illustrates the generated intron-less strains expressing either Heh1-L or Heh1-S. Asterisk denotes premature splice recognition site. Right panel: Immunoblotting of Heh1-L^6HA^, Heh1-S^6HA^ and Hub1 from cells grown in YPD at 30°C and shifted to 23°C, 30°C or 37°C for 3h (asterisk denotes a cross-reacting signal). Dpm1 was used as a loading control. (D,E) Five-fold serial dilutions of the indicated strains were spotted and grown on YPD at 30°C or 37°C for 3 days. (F) Left panel: Representative pictures of live-cell imaging of Nup192^GFP^ in the indicated yeast strains (arrows indicate SINCs). Maximum intensity Z-projections are shown. Right panel: Quantification of the percentage of cells that accumulate Nup192^GFP^ at the SINC (highlighted with arrows) in the indicated strains. Error bars represent ± SD from two independent replicates (n > 1000 cells for each strain). Analysis of variance (ANOVA) was performed, and different letters denote significant differences with a Tukey’s *post hoc* test at P < 0.05.

### Heh1-S is essential for normal growth of ESCRT-III mutants

Hub1 promotes the generation of the shorter spliced variant Heh1-S, while its absence causes nearly exclusive expression of the long form Heh1-L (Karaduman et al., 2017; Mishra et al., 2011; Fig. S1A,B). To discern whether the synthetic growth defect of *hub1*Δ *did4*Δ cells is due to defective *HEH1* splicing, we generated yeast strains that exclusively express either Heh1-L or Heh1-S from the endogenous locus (*HEH1-L* and *HEH1-S* strains, respectively; Fig. 1C). Both intron-less variants display similar expression levels and nuclear localization (Fig. 1C; Fig. S1C). Using a functional assay for NPC integrity (Yewdell et al., 2011), we further confirmed that both alleles fully complement the *heh1*Δ phenotype (Fig. S1D,E). These results suggest that both spliced versions are functional and act redundantly in proper NPC assembly.

Next, we used the *HEH1-L* and *HEH1-S* strains to study whether concomitant deletion of *HEH1-S* affects growth in strains lacking ESCRT proteins. Similar to the *hub1*Δ mutant (Fig. 1A,B), the additional absence of Heh1-S produced a severe growth defect in *did4*Δ or *did2*Δ cells (ESCRT-III mutants), whereas removal of Heh1-L caused no detectable phenotype (Fig. 1D,E; Fig. S2A,B). Similarly, combining *HEH1-L* with mutants lacking ESCRT-0, ESCRT-I, ESCRT-II or ESCRT accessory genes did not aggravate the growth defect (Fig. S2), suggesting that this genetic interaction is specific for ESCRT-III.

Deletion of ESCRT-III components results in clustering of NPCs at the SINC due to disruption of the pathway that removes defective NPC assembly intermediates (Webster et al., 2014, 2016). To assess whether lack of Heh1-S affects NPC clustering, we analyzed SINC formation in *did4*Δ and *HEH1-L did4*Δ strains using a C-terminal GFP fusion of the nucleoporin Nup192 (Nup192^GFP^). In agreement with previous studies (Webster et al., 2014, 2016), we found a significant increase of Nup192 accumulation at the SINC in the ESCRT-III mutant compared to WT and *HEH1-L* cells (10% vs. 3-4%; Fig. 1F). However, Nup192 accumulation was further increased in *did4*Δ cells expressing only Heh1-L (more than 30%; Fig. 1F). Together, these results imply that the exclusive presence of Heh1-L contributes to the accumulation of NPCs at the SINC in cells deficient in the ESCRT-III machinery.

### Phenotypes associated with Heh1-S deficiency depend on the MSC domain of Heh1-L

Heh1-L contains two conserved domains exposed to the nucleoplasm: the LEM and the MSC domain (Fig. 1C, left panel). In *Schizosaccharomyces pombe*, these domains perform distinct functions in the Heh1-L homolog, Lem2 (Barrales et al., 2016). We sought to investigate which domain of Heh1 is responsible for the synthetic growth defect of ESCRT-III mutants expressing only Heh1-L. To this end, we generated multiple truncated versions of Heh1 lacking individual nucleoplasmic or transmembrane domains (Δ*LEM*, Δ*TM1*, Δ*TM2*, Δ*MSC)* or combinations thereof (Δ*TM1-TM2* and Δ*TM2-MSC*) and expressed them as plasmid-borne N-terminal GFP fusions under the control of the inducible *GAL1* promoter in strains lacking endogenous *HEH1* (Fig. 2A). Most of the truncated versions were expressed at similar levels as Heh1-L and localized to the NE, with the exception of those lacking the first transmembrane domain, which displayed reduced expression and had mainly nucleoplasmic localization (Fig. S3A,B). While *heh1*Δ *did4*Δ cells display normal growth, overexpression of full-length *HEH1-L* causes a severe growth defect. Notably, deleting the MSC domain nearly completely rescued this growth defect (Fig. 2B). A comparable outcome was found for constructs in which the MSC domain is exposed to the lumen of the NE or Heh1-L is not membrane-bound (Δ*TM1* and Δ*TM1-TM2*; Fig. 2B). In contrast, lack of the LEM domain did not suppress the growth defect (Fig. 2B). Similar results were obtained for *heh1*Δ *did4*Δ and *heh1*Δ *did2*Δ cells grown at 37°C, although under this condition the suppression of the Heh1-L-mediated phenotype was partially weaker (Fig. S3C,D). In *S. pombe*, a region of Lem2 adjacent to the first TM mediates interaction with another nuclear membrane protein, Bqt4 (Hirano et al., 2018; Hu et al., 2019). However, expression of mutants lacking the equivalent region in Heh1-L or other N-terminal regions affected the growth of *heh1*Δ *did4*Δ cells to the same extent or even stronger as the full-length protein (Fig. S3E). These findings suggest that the toxicity of Heh1-L does not require N-terminal domains. Recently, it was shown that cytosolic exposure of the MSC domain of Heh1-L is sufficient to recruit the ESCRT machinery to the NE (Thaller et al., 2019). We therefore assessed the phenotype of two distinct Heh1-L mutants that lack a functional <u>n</u>uclear localization <u>s</u>ignal (NLS), resulting in the cytoplasmic exposure of the MSC domain (Δ*NLS* and *R176A*; Lokareddy et al., 2015; Fig. 2A). Notably, these mutants display an even more severe growth defect than WT Heh1-L when expressed in *heh1*Δ *did4*Δ cells (Fig. 2C; Fig. S3F). Together, these findings imply that the MSC domain is responsible for the slow growth phenotype of ESCRT-III mutants expressing only Heh1-L.

**Fig. 2.**
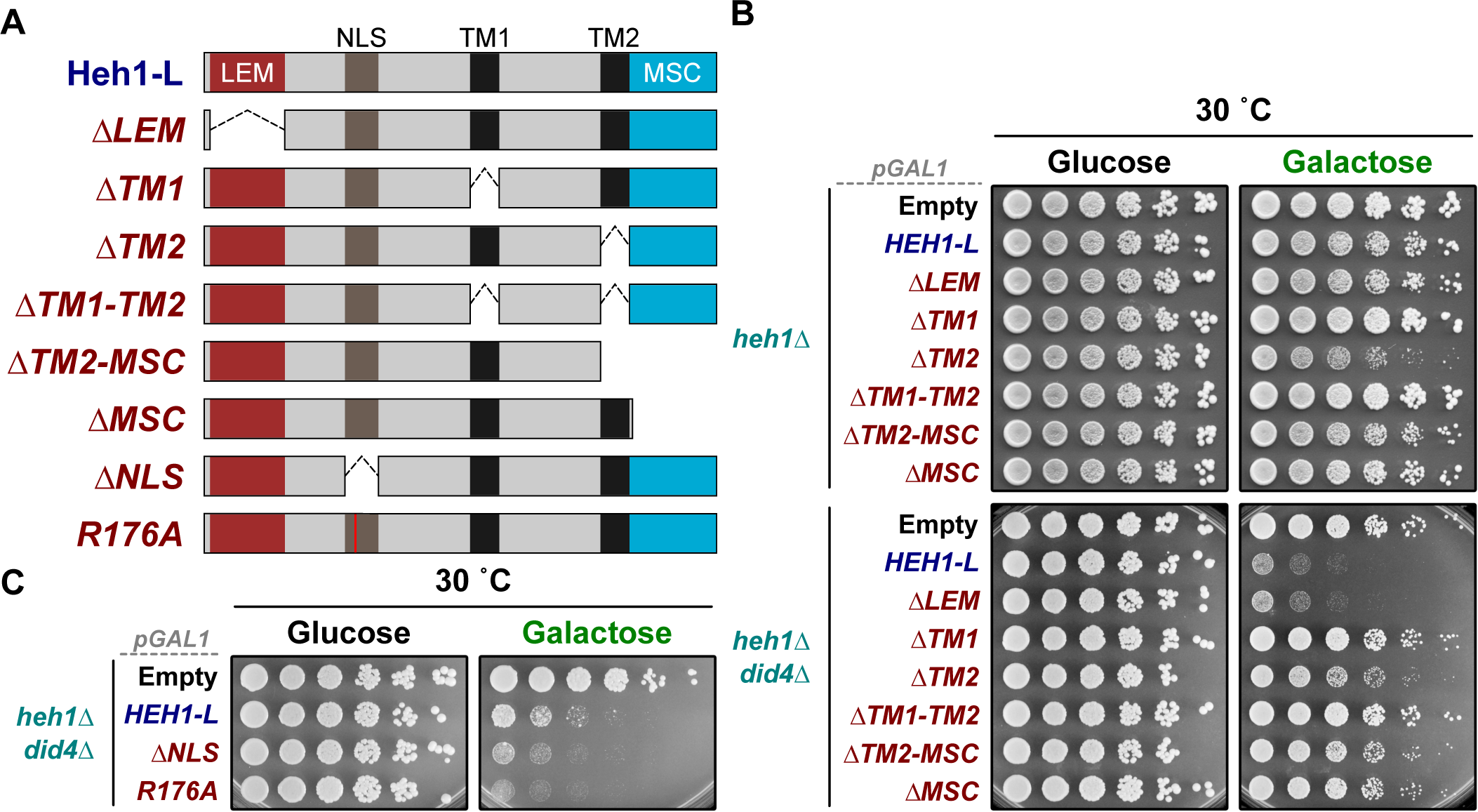
The MSC domain of Heh1-L is critical for the growth defect of ESCRT-III mutants. (A) Scheme illustrating the truncated constructs of Heh1-L. (B) Five-fold serial dilutions of the indicated strains were spotted and grown on selective media with glucose (control) or galactose (induction) at 30°C for 3 days. The different strains were transformed with empty vector or plasmids bearing GFP-tagged *HEH1-L* or the different truncated mutants with the galactose-inducible promoter. (C) Five-fold serial dilutions of *heh1*Δ *did4*Δ strains were spotted and grown on selective media with glucose (control) or galactose (induction) at 30°C for 3 days. The different strains were transformed with empty vector or plasmids bearing GFP-tagged *HEH1-L* or the different NLS mutants with the galactose-inducible promoter.

LEM-containing proteins are highly conserved from yeast to humans (Brachner and Foisner, 2011). To identify conserved residues within the MSC domain that are potentially critical for its function, we performed sequence alignment of proteins from different species. *In silico* analysis of this region revealed several conserved residues (Fig. 3A), which we mutated into alanine (A) and assessed their functional relevance. Using immunoblotting and live-cell imaging, we verified the proper expression and localization of these MSC mutants (Fig. 3B; Fig. S4A,B). Most of them also displayed a growth phenotype similar to WT Heh1-L when expressed in *heh1*Δ *did4*Δ cells. However, specific tryptophan (W) to A mutants were found to partially suppress the phenotype of Heh1-L, especially when combined with each other (e.g. W786A and W817A; Fig. 3C; Fig. S4C). Interestingly, those conserved tryptophan residues form a hydrophobic surface in the MSC of human MAN1 but are not present in other winged-helix domains (Caputo et al., 2006). From these data we conclude that the presence of a functional MSC domain of Heh1-L triggers the growth defect observed in ESCRT-III mutant cells.

**Fig. 3.**
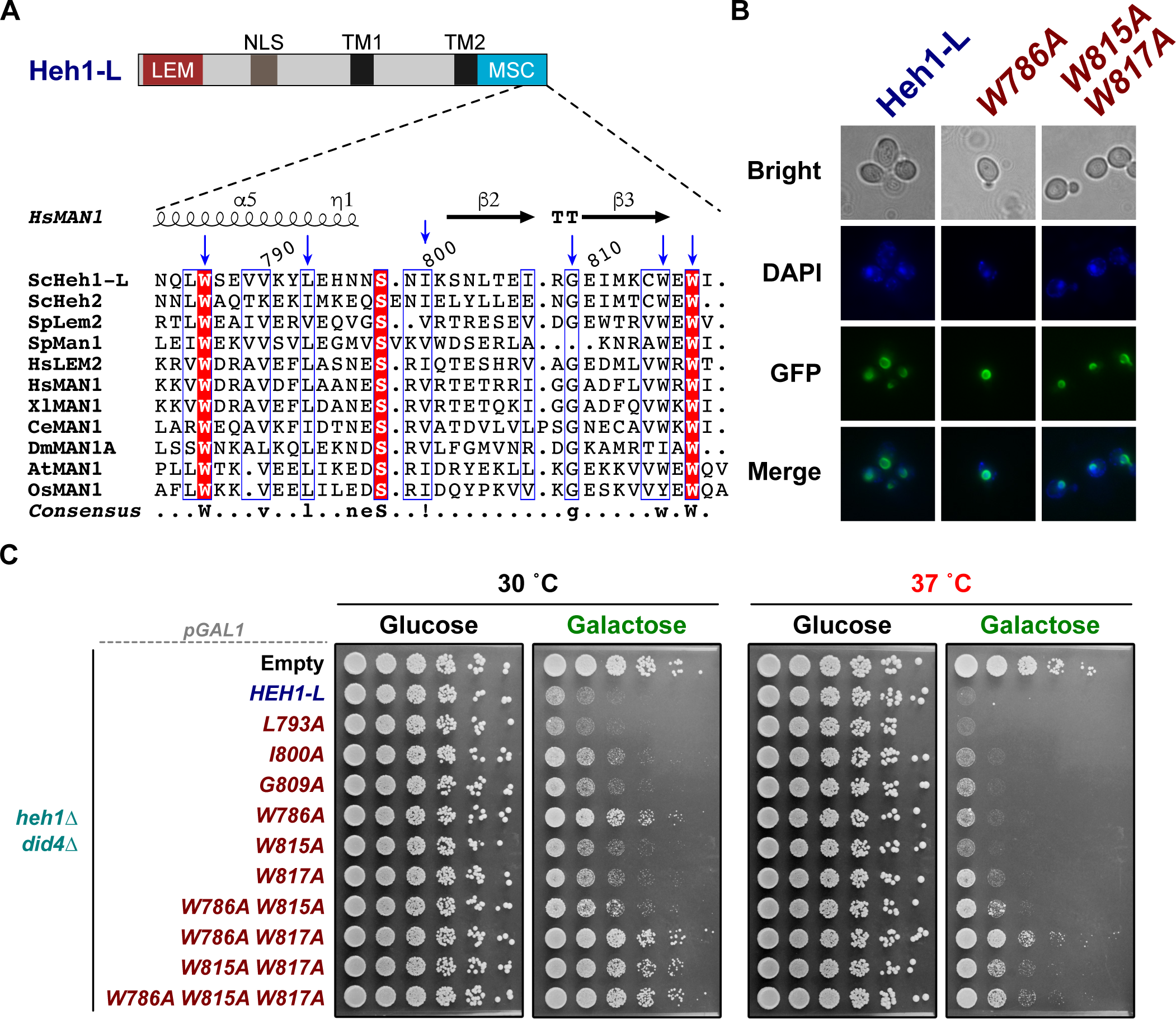
Conserved residues at the MSC domain of Heh1-L are involved in mediating the growth defect of ESCRT-III mutants. (A) Sequence alignment and secondary structure elements of the MSC domains from *Saccharomyces cerevisiae* (Sc), *Schizosaccharomyces pombe* (Sp), *Homo sapiens* (Hs), *Xenopus laevis* (Xl), *Caenorhabditis elegans* (Ce), *Drosophila melanogaster* (Dm), *Arabidopsis thaliana* (At), and *Oriza sativa* (Os). The secondary structure elements are based on the crystal structure of human MAN1 and are shown above the sequences (α = alpha helix, ß = beta strand, η = 3_10_ helix). Aligned residues in white with red highlight are conserved among species, and blue boxes denote conserved cluster of amino acids. The mutated residues are marked with a blue arrow. (B) Live-cell imaging of *heh1*Δ cells transformed with plasmids bearing ^*GFP*^*HEH1-L* or the point mutants ^*GFP*^*heh1-L W786A* and ^*GFP*^*heh1-L W815A W817A* with the galactose-inducible promoter after 2h of induction. DAPI staining was used as a nuclear marker. An individual image of one of the Z-stacks is shown. (C) Five-fold serial dilutions of the indicated strains were spotted and grown on selective media with glucose (control) or galactose (induction) at 30°C for 3 days. The different strains were transformed with empty vector or plasmids bearing ^*GFP*^*HEH1-L* or the different point mutants with the galactose-inducible promoter.

### The growth defect of cells lacking Heh1-S and ESCRT-III is caused by Chm7

Heh1-L recruits the ESCRT machinery to the NE through interaction with the chimeric ESCRT-II/III protein Chm7 (Bauer et al, 2015; Gu et al., 2017; Thaller et al., 2019; Webster et al., 2016). Chm7 exists in an inactive or active (open state) conformation due to the presence of auto-inhibitory helices (Lata et al., 2008; Webster et al., 2016). Recruitment to the NE likely requires Chm7 being present in the open state (Thaller et al., 2019). In other species, recruitment of Chm7 homologs further requires the presence of the MSC domain of Lem2, which is Heh1-L homolog (Gu et al., 2017; Pieper et al., 2020; Thaller et al., 2019; von Appen et al., 2020). We therefore assessed whether the constitutively active version of Chm7 (chm7_OPEN_) is critical for binding to the MSC domain of Heh1-L. Using yeast two-hybrid (Y2H) and co-immunoprecipitation (coIP) assays, we found that the MSC domain of Heh1-L interacts with chm7_OPEN_ but not full-length Chm7 (Fig. 4A,B; Fig. S5A). Moreover, as expected, Heh1-S that lacks the MSC domain showed no binding to Chm7 (Fig. 4B; Fig. S5A). To test whether the residues in Heh1-L that contribute to its toxicity when overexpressed in ESCRT-III mutants are also critical for interaction with Chm7, we examined those mutants on their binding to chm7_OPEN_. Indeed, W786A and W817A mutations diminished the binding of Chm7 to the MSC of Heh1-L (Fig. 4C,D). We further found that a mutation in which the MSC is facing the NE lumen (Δ*TM2*) completely abolished the interaction (Fig. 4D). These results demonstrate that the MSC domain preferentially binds to chm7_OPEN_ and that this interaction involves the residues W786 and W817 of Heh1-L.

**Fig. 4.**
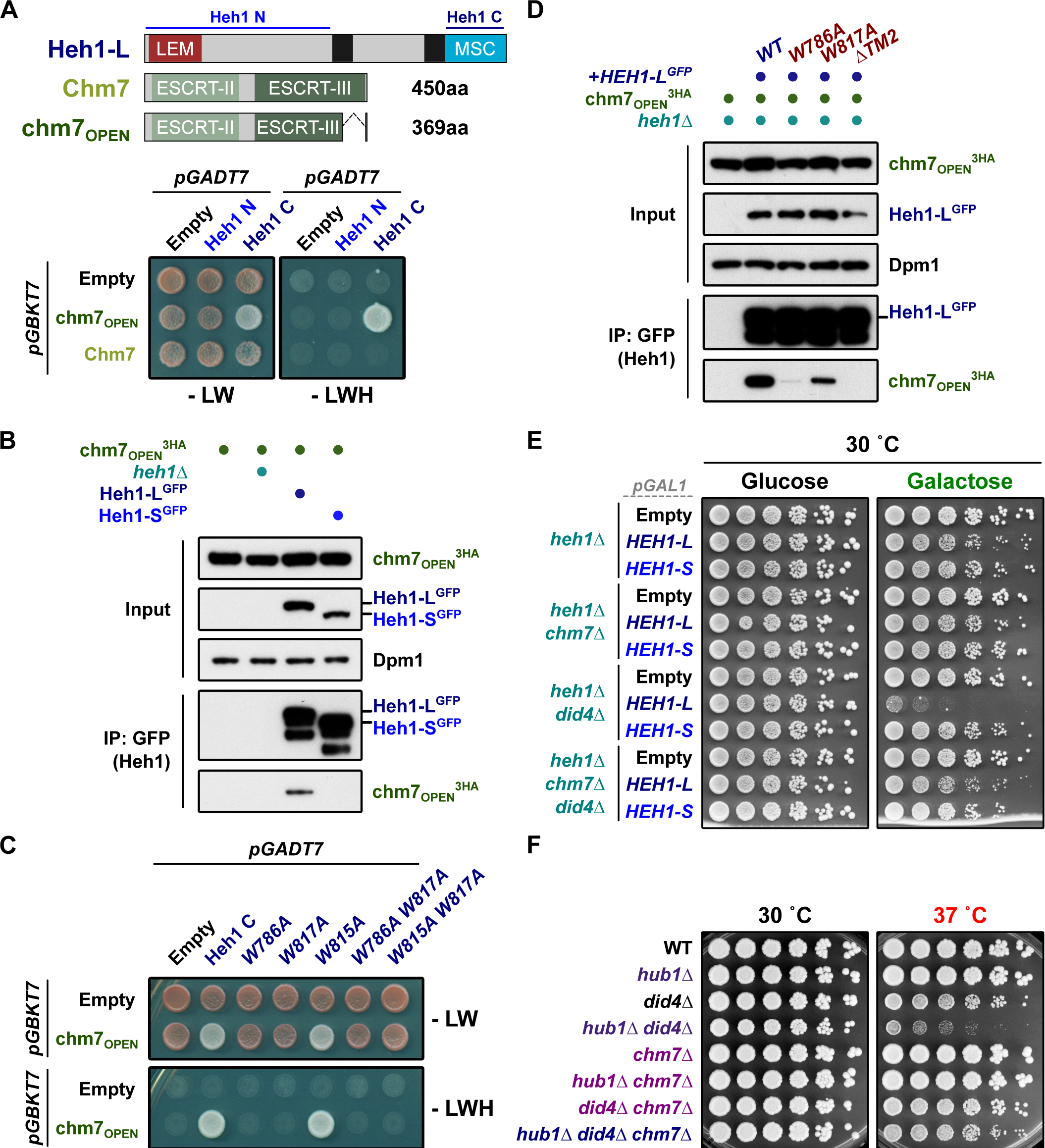
The MSC domain of Heh1-L recruits chm7_OPEN_, causing growth defects in ESCRT-III mutants. (A) Upper panel: schematic representation of the constructs used in yeast two-hybrid (Y2H) assays. The protein length of Chm7 and chm7_OPEN_ is indicated. Lower panel: Y2H analysis of Chm7 or chm7_OPEN_ (as a Gal4-DNA-binding domain fusion in the *pGBKT7* vector) with either Heh1 N or Heh1 C (as a Gal4-activating domain fusion in the *pGADT7* vector). (B) Interaction of chm7_OPEN_^3HA^ with either Heh1-L^GFP^ or Heh1-S^GFP^ was detected by co-immunoprecipitation. Dpm1 served as a loading control. (C) Y2H analysis of chm7_OPEN_ (as a Gal4-DNA-binding domain fusion in the *pGBKT7* vector) with Heh1 C and several point mutants as indicated (as a Gal4-activating domain fusion in the *pGADT7* vector). (D) Interaction of chm7_OPEN_^3HA^ with Heh1-L^GFP^ or the *heh1-L*^*GFP*^ *W786A, heh1-L*^*GFP*^ *W817A* or *heh1-L*^*GFP*^ Δ*TM2* mutants was detected by co-immunoprecipitation. Dpm1 served as a loading control. The chm7_OPEN_^3HA^ *heh1*Δ strain was transformed with empty vector or plasmids bearing the different *HEH1-L* constructs with the endogenous promoter (indicated with +). (E) Five-fold serial dilutions of the indicated strains were spotted and grown on selective media with glucose (control) or galactose (induction) at 30°C for 3 days. The different strains were transformed with empty vector or plasmids bearing GFP-tagged *HEH1-L* or *HEH1-S* with the galactose-inducible promoter. (F) Five-fold serial dilutions of the indicated strains were spotted and grown on YPD at 30°C or 37°C for 3 days. For A and C, cells were spotted on control media (-LW) or selective media (-LWH) and grown for 3 days.

Since Chm7 is required for SINC formation and interacts with Heh1-L (Thaller et al., 2019; Webster et al., 2016), we speculated that Chm7 might directly contribute to the growth phenotype of ESCRT-III mutants lacking Heh1-S. To test this hypothesis, we examined the growth of cells expressing exclusively Heh1-L in ESCRT-III mutants (either *did4*Δ or *did2*Δ) in presence and absence of Chm7. Remarkably, the severe growth phenotype of Heh1-L overexpression was nearly fully suppressed when Chm7 was absent (Fig. 4E; Fig. S5B-D). Consistent with our findings, overexpression of Heh1-S did not cause any growth defect (Fig. 4E, Fig. S5B-D). Finally, we tested whether the synthetic interaction of *HUB1* and *DID4* also depends on *CHM7*. Indeed, deleting *CHM7* in cells lacking Hub1 and Did4, completely rescued the growth phenotype (Fig. 4F). Altogether, these data suggest that uncontrolled association between Chm7 and Heh1-L is detrimental in cells lacking the ESCRT-III complex.

### Heh1-S impairs the interaction between Chm7 and Heh1-L

Encouraged by our finding that Heh1-L binds specifically to the open conformation of Chm7, we wondered whether exclusive expression of Heh1-L in *chm7*_*OPEN*_ cells results in a growth defect analogous to mutants lacking ESCRT-III members (Fig. 1D,E). While we observed no defect under normal conditions (Fig. 5A, left panel), we found that the sole presence of Heh1-L, but not Heh1-S, caused significant sensitivity towards DMSO (Fig. 5A, right panel), which has been reported to affect the structure and properties of biological membranes (Gurtovenko and Anwar, 2007; Notman et al., 2006). Conversely, deletion of *CHM7* improved growth in the presence of DMSO (Fig. 5A). Intriguingly, the severe growth phenotype upon DMSO treatment of *HEH1-L chm7*_*OPEN*_ cells was rescued when plasmid-borne *HEH1-S* was concomitantly expressed from its endogenous promoter (Fig. 5B). We then explored the mechanism by which Heh1-S rescues the growth defect of *HEH1-L* cells expressing *chm7*_*OPEN*_ or lacking ESCRT-III members. By performing coIP experiments in the presence or absence of Heh1-S, we found that co-expression of Heh1-S weakens the interaction between Chm7 and Heh1-L (Fig. 5C, compare lanes 2 and 3). Together, these findings imply that Heh1-S interferes with the binding of Chm7 to Heh1-L and prevents the toxicity of constitutively active Chm7, especially under conditions that affect membrane organization.

**Fig. 5.**
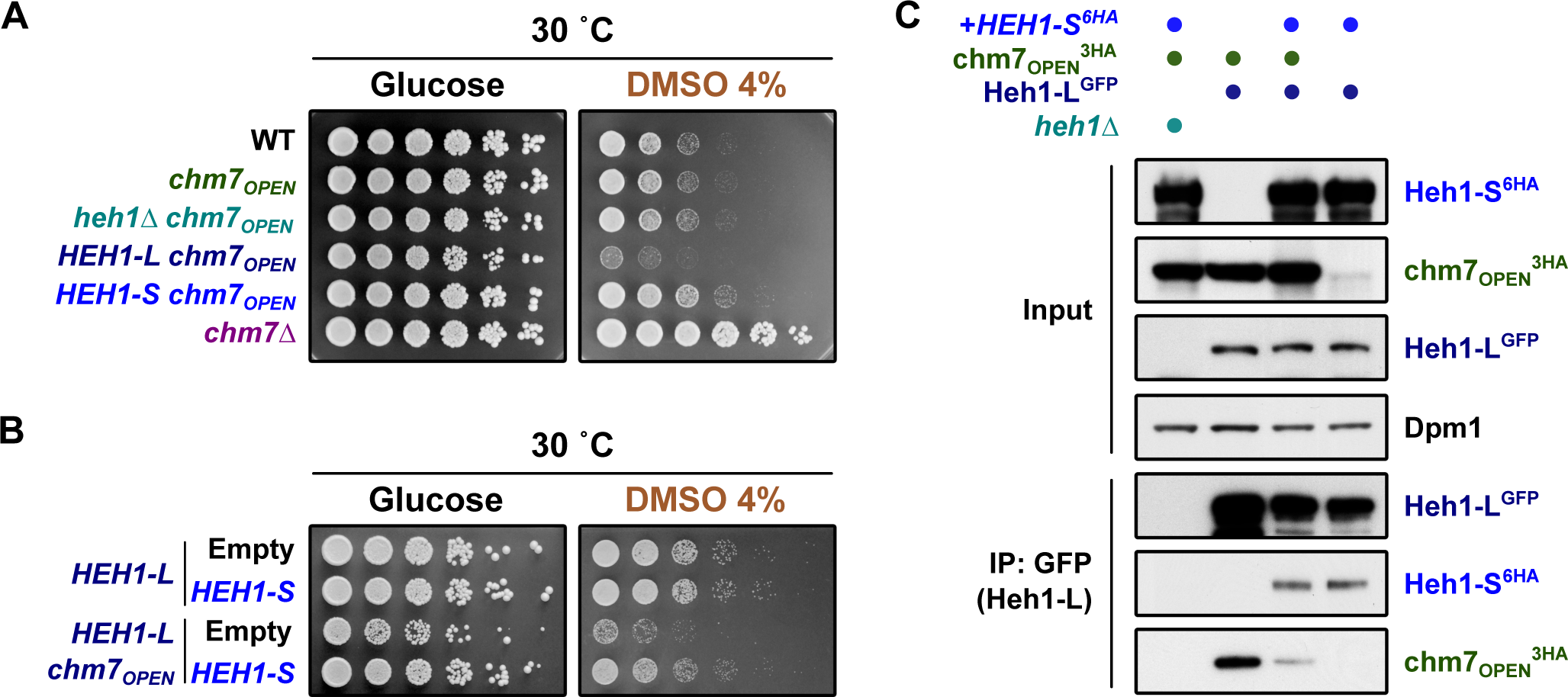
The presence of Heh1-S interferes with Chm7 binding to Heh1-L. (A) Five-fold serial dilutions of the indicated strains were spotted on minimal media plates, with or without DMSO 4%, and were incubated for 3 days at 30°C. (B) Five-fold serial dilutions of the indicated strains were spotted and grown on selective media, with or without DMSO 4%, and were incubated for 3 days at 30°C. The different strains were transformed with empty vector or plasmids bearing GFP-tagged *HEH1-S* with the endogenous promoter. (C) Interaction of Heh1-L^GFP^ with either chm7_OPEN_^3HA^ or Heh1-S^6HA^ was detected by co-immunoprecipitation. The different strains were transformed with empty vector or a plasmid bearing GFP-tagged *HEH1-S* with the endogenous promoter (indicated by +).

### A unique sequence element of Heh1-S is critical to control the interaction of Heh1-L and Chm7

In higher eukaryotes, LEM-containing proteins have been shown to oligomerize (Berk et al., 2014; Mansharamani and Wilson, 2005). We therefore examined whether Heh1-S can bind to Heh1-L, which might explain its ability to prevent the interaction of Heh1-L with Chm7. By expressing different epitope-tagged versions of Heh1-L and Heh1-S, we found that both of them can form homo- and heterodimers (Fig. S6A). Using Y2H experiments, we then mapped the interaction between Heh1-L and Heh1-S to individual domains (Fig. 6A,B). We observed that the heterodimerization interface comprises a region of the LD (lumen domain) shared between both forms (residues 476 to 638; Fig. 6B; Fig. S6B). This result was further confirmed by coIP (Fig. 6C). Together, these findings demonstrate that Heh1-S and Heh1-L interact through their lumen domain.

**Fig. 6.**
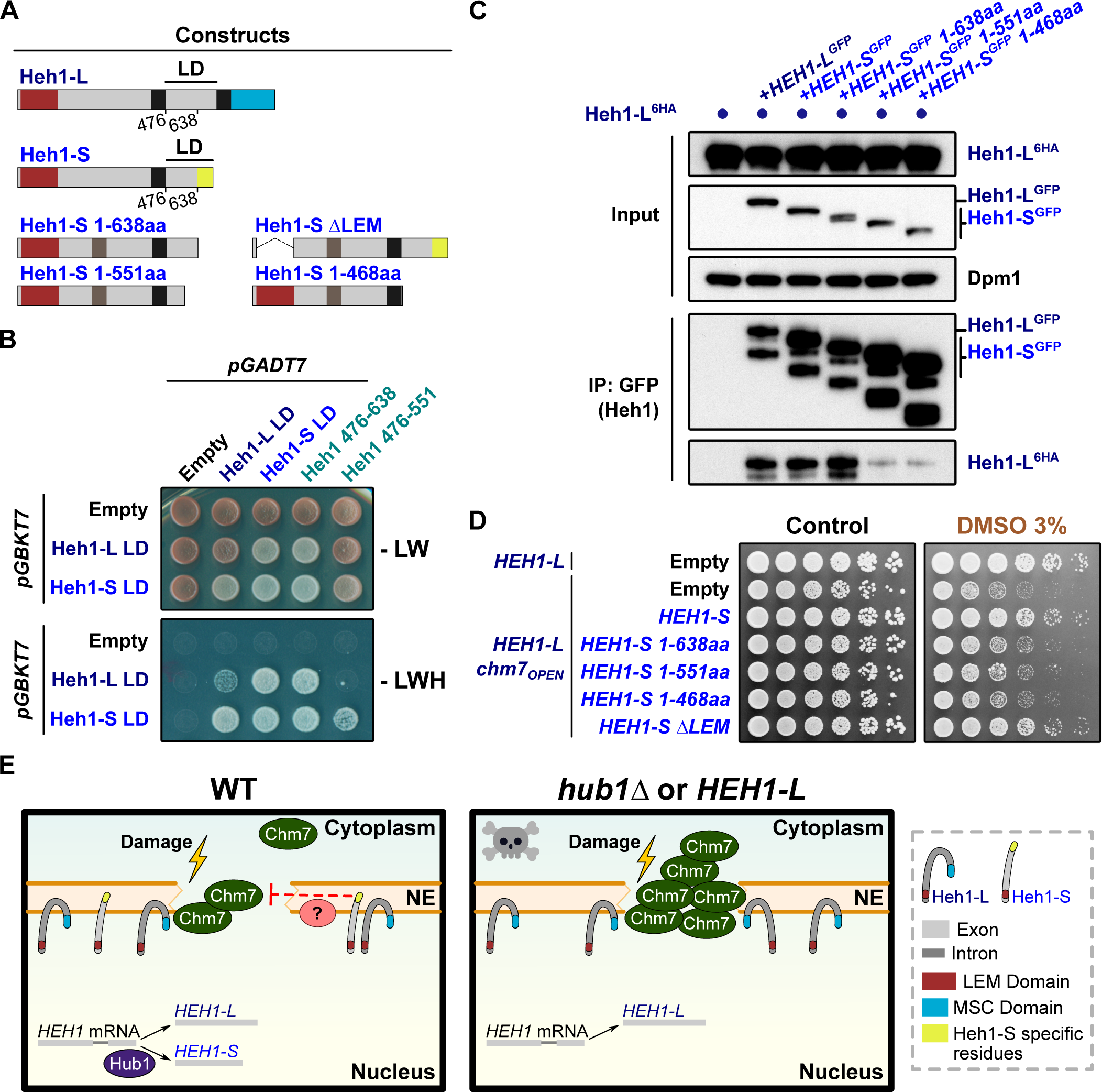
Unique residues in Heh1-S are required to rescue the growth defect of *HEH1-L chm7*_*OPEN*_ cells. (A) Schematic representation of the constructs used for Y2H, co-immunoprecipitation and complementation assays. (B) Y2H analysis of the lumen domain (LD) of Heh1-L and Heh1-S (as a Gal4-DNA-binding domain in the *pGBKT7* vector, and Gal4-activating domain fusions, in the *pGADT7* vector) and two truncated mutants as indicated (as a Gal4-activating domain fusion in the *pGADT7* vector). Cells were spotted on control media (-LW) or selective media (-LWH) and grown for 3 days at 30°C. (C) Interaction of Heh1-L^6HA^ with Heh1-L^GFP^, Heh1-S^GFP^ or the truncated mutants of Heh1-S^GFP^ (1-638aa, 1-551aa and 1-468aa) was detected by co-immunoprecipitation. C-terminally GFP-tagged constructs were expressed from plasmids under the control of *HEH1* endogenous promoter (indicated by +). (D) Five-fold serial dilutions of *HEH1-L* and *HEH1-L chm7*_*OPEN*_ strains were spotted and grown on selective media, with or without DMSO 3%, and were incubated for 3 days at 30°C. The different strains were transformed with empty vector or plasmids bearing GFP-tagged *HEH1-S* (WT or truncated mutants) with the endogenous promoter. (E) Proposed model of Heh1-S function at the NE. In WT cells, Hub1-mediated alternative splicing allows the expression of Heh1-L and Heh1-S. When perturbations occur at the NE (indicated by lightning symbol), Heh1-L recruits Chm7 to mediate repair and this interaction is modulated by the presence of Heh1-S (left panel), which could be potentially assisted by yet-unknown factor/s (?). In cells lacking Heh1-S (*hub1*Δ or *HEH1-L* cells), Chm7 is excessively recruited to the NE (right panel) giving rise to a growth defect. NE, nuclear envelope.

We next explored whether Heh1-S directly competes with Chm7 for Heh1-L association; however, we observed no difference in Heh1-L and Heh1-S heterodimer formation in the presence of chm7_OPEN_ (Fig. 4C; compare co-immunoprecipitated Heh1-S^6HA^ in lane 3 and 4). Furthermore, Heh1-L requires its MSC domain to bind Chm7 (Fig. 4A,C), whereas the LD is needed for Heh1-S interaction (Fig. 6B). This suggests that Heh1-S presents a distinct feature that is necessary to impair Chm7 recruitment to the NE. Indeed, Heh1-S contains 49 unique residues at its C-terminal (Grund et al., 2008; Rodríguez-Navarro et al., 2002), which present a high degree of conservation among yeast species (Fig. S6E). Notably, we found that a Heh1-S mutant lacking these unique residues was unable to suppress the sensitivity of *HEH1-L chm7*_*OPEN*_ cells towards DMSO (Heh1-S 1-638aa; Fig. 6D). Interestingly, this unique fragment was not required for heterodimerization (Fig. 6C, compare lane 4 with lanes 5 and 6). Conversely, complementation was independent of the LEM domain (Δ*LEM*; Fig. 6D). We could exclude that lack of complementation was due to altered protein stability, since all examined mutants were expressed to similar levels compared to full-length Heh1-S (Fig. S6C). Furthermore, expression of N-terminal truncations of Heh1-S also rescued the growth defect of *HEH1-L chm7*_*OPEN*_ cells (Fig. S6D). Together, these results suggest that the unique residues of Heh1-S modulate the interaction between Heh1-L and Chm7, and imply that suppression of the growth defect cannot be explained only by heterodimerization. We thus conclude that the spliced form Heh1-S modulates the interaction of Heh1-L with Chm7 and prevent its excessive recruitment to the nuclear envelope.

## Discussion

The ubiquitin-relative Hub1 assists pre-mRNA splicing in yeast and human cells (Ammon et al., 2014; Mishra et al., 2011; Wilkinson et al., 2004). Although Hub1 up-regulation is correlated with cadmium tolerance in budding yeast (Chanarat and Svasti, 2020), the physiological role of Hub1-mediated splicing remains poorly understood. Here, we unveil that Hub1 modulates the ESCRT recruitment to the NE by regulating the alternative splicing of the inner nuclear membrane protein Heh1.

Whereas Heh1-L recruits the ESCRT member Chm7 to seal NE holes, persistent interaction between them is toxic for cells (Thaller et al., 2019; Webster et al., 2016; Willan et al., 2019), indicating that this process needs to be tightly regulated. This association is resolved by the ESCRT-III/Vps4 complex, since depletion of ESCRT-III members causes Chm7 accumulation at the NE in yeast and human cells (Gu et al., 2017; Pieper et al., 2020; Webster et al., 2016; Willan et al., 2019). In addition, lack of the AAA ATPase Vps4 in fission yeast causes a growth defect accompanied by NE defects, which can be suppressed by deleting Chm7 or blocking its recruitment to the NE (Gu et al., 2017; Pieper et al., 2020). In agreement with these findings, we provide several lines of evidence that uncontrolled Heh1-L-mediated recruitment of Chm7 is the underlying cause of the growth defect in ESCRT-III mutants. First, masking or complete removal of the MSC domain suppresses the synthetic growth phenotype of Heh1-L overexpression in ESCRT-III mutants (Fig 2B; Fig. S3). Second, a similar outcome is seen when Heh1-L is expressed with mutations at its MSC domain that specifically disrupt the interaction with Chm7 (Figs. 3C, 4D). Lastly, this synthetic growth phenotype strictly depends on the presence of Chm7 (Fig. 4E).

In *S. cerevisiae*, Chm7 is constitutively exported from the nucleus by the major exportin Crm1, suggesting the existence of an active mechanism to physically separate Chm7 and Heh1-L (Thaller et al., 2019). In agreement with this finding, we found that overexpression of nuclear import-deficient Heh1-L mutants in the ESCRT-III *did4*Δ mutant caused a stronger phenotype than expression of WT Heh1-L (Fig. 2C; Fig. S3). Moreover, CHMP7, the human homolog of Chm7, also contains a functional nuclear export signal, indicating a similar regulation in mammalian cells (Vietri et al., 2019 preprint). Our findings imply an additional layer of regulation that involves the alternative splicing of Heh1. Cells lacking Hub1, which are compromised in producing the spliced variant Heh1-S, display impaired growth when combined with ESCRT-III mutants (*did2*Δ or *did4*Δ) and accumulate misassembled NPCs at the SINC (Fig. 1; Fig. S2). Importantly, this phenotype seems to arise from a toxic gain-of-function of Heh1-L and not to the mere absence of Heh1-S, since no synthetic growth phenotype was detected in *heh1*Δ cells. Furthermore, deletion of *CHM7* also rescued the synthetic growth phenotype of *hub1*Δ *did4*Δ cells (Fig. 4F), further supporting the idea that this defect is due to the lack of *HEH1* splicing. Importantly, we found that the interaction between Chm7 and Heh1-L is impaired by Heh1-S expression (Fig. 5C), implying that it prevents or counteracts excessive Chm7 recruitment. Intriguingly, this behavior is highly reminiscent of the regulation of human Siah-1 (<u>s</u>even <u>i</u>n <u>a</u>bsentia <u>h</u>omolog), an ubiquitin ligase implicated in several cellular functions. Alternative splicing generates the shorter variant Siah-1S. The short version forms heterodimers with Siah-1, which seems to be critical for complex formation with Siah-interacting protein (Mei et al., 2007).

A recent study showed that human LEM2 condensates into a liquid-like phase through intrinsically disordered regions (IDR) present at its N-terminal domain, which seem to be critical during NE reassembly and postmitotic nucleocytoplasmic compartmentalization (von Appen et al., 2020). Interestingly, Heh1 also has IDRs at its N-terminal domain (Fig. S7), suggesting that a similar process may also take place in yeast. However, overexpression of Heh1-L lacking the LEM domain or the N-terminal region impairs growth of ESCRT-III mutants to the same extent as the full-length protein (Fig. 2B; Fig. S3C-E). Conversely, the growth defect of *HEH1-L chm7*_*OPEN*_ cells can be rescued by the expression of LEM- or N-terminal truncated versions of Heh1-S (Fig. 6D; Fig. S6D). While this does not exclude the possibility that other functions of Heh1 may be influenced by the presence of the N-terminal IDRs, these data suggest that at least Chm7 recruitment and its regulation by Heh1-S is controlled in a manner independent of N-terminal motifs.

The specific role of Heh1-S in modulating the recruitment of ESCRT-III to the NE implies the presence of distinct features not present in Heh1-L. Besides the absence of the second transmembrane domain and the MSC domain, Heh1-S contains 49 unique residues at its C-terminus due to the frameshift generated by Hub1-mediated alternative splicing of *HEH1* (Grund et al., 2008; Mishra et al., 2011; Rodríguez-Navarro et al., 2002). Interestingly, while not involved in heterodimerization with Heh1-L, the presence of these residues in Heh1-S is critical to rescue DMSO toxicity in *HEH1-L chm7*_*OPEN*_ cells (Fig. 6D). Hence, we propose a model in which the repair of membrane damage at the NE is fine-tuned by the two splice products of *HEH1*. NE ruptures or misassembled NPCs promote the interaction between Heh1-L and Chm7, which in turn recruits ESCRT-III proteins and Vps4 to repair damage through membrane scission. In wild-type cells, Hub1-spliced Heh1-S modulates the interaction between Chm7 and Heh1-L (Fig. 6E, left panel). The mechanism by which Heh1-S regulates the recruitment of Chm7 still remains elusive. We speculate that the unique residues of Heh1-S might recruit a yet-unidentified factor to avoid excessive NE remodeling. However, in cells lacking Heh1-S, Chm7 recruitment is not regulated (Fig. 6E, right panel), leading to its continuous recruitment. This most likely results in excessive activity and deleterious phenotypes, such as membrane fenestration or “karmallae” formation, as reported in other species (Gu et al., 2017).

Our data provide a novel example for the key role that alternative splicing plays in regulating a specific cellular function, such as NE integrity. Given the strong conservation of LEM domain proteins, our results may provide insights into an ancient molecular mechanism evolved to avoid excessive NE remodeling.

## Materials and methods

### Yeast methods and molecular biology

*S. cerevisiae* strains and plasmids are listed in Tables S1 and S2, respectively. Standard protocols for transformation, mating, sporulation and tetrad dissection were used for yeast manipulations. Yeast growth assays were performed by spotting five-fold serial dilutions of the indicated strains on solid agar plates. Yeast strains isogenic to W303 or DF5 were used for genetic studies, and PJ69-7a for yeast two-hybrid assays. Cells were grown at 30°C (if not indicated otherwise) in YPD medium (2% glucose) or synthetic complete medium lacking individual amino acids to maintain selection for transformed plasmids. Protein tagging and the construction of deletion mutants were conducted by a PCR-based strategy (Janke et al., 2004; Knop et al., 1999) and confirmed by immunoblotting and PCR, respectively. For galactose induction, cells were cultured in synthetic complete medium lacking the specific amino acids + 2% raffinose, and galactose was added to a mid-log phase yeast cells to a final concentration of 2%. Cloning methods used standard protocols or the Gibson Assembly Master Mix (NEB). *HEH1* and *CHM7* were cloned into *pGADT7* or *pGBKT7* vectors (Clontech™) for yeast two-hybrid experiments. Plasmids with point mutations or deletions were constructed by the PCR-based site-directed mutagenesis approach. Maps and primer DNA sequences are available upon request.

### Immunoblotting

Total protein extracts from 2×10^7^ cells (OD_600_=1) were prepared by trichloroacetic acid (TCA) precipitation (Knop et al., 1999). Proteins solubilized in HU loading buffer (8 M urea, 5% SDS, 200 mM Tris-HCl pH 6.8, 20 mM dithiothreitol (DTT) and bromophenol blue 1.5 mM) were resolved on NuPAGE 4%-12% gradient gels (Invitrogen), and analyzed by standard immunoblotting techniques. Dpm1 antibody was used as loading control.

### Antibodies

Polyclonal Hub1 antibodies were raised in rabbits and have been described previously (Mishra et al., 2011). Monoclonal (3F10) antibody directed against the HA epitope and monoclonal (B-2) antibody against GFP were purchased from Roche and Santa Cruz Biotechnology (Dallas, TX), respectively. Monoclonal Dpm1 (5C5) antibody was purchased from Thermo Fisher Scientific.

### Live-cell imaging and analysis

Imaging was performed on a Zeiss AxioObserver Z1 confocal spinning-disk microscope equipped with an Evolve 512 (Photometrics) EMM-CCD camera through a Zeiss Alpha Plan/Apo 100x/1.46 oil DIC M27 objective lens. Optical section images were obtained at focus intervals of 0.2 μm. Subsequent processing and analyses of the images were performed in Fiji/ImageJ (Schindelin et al., 2012). DAPI staining was performed by adding 2.5 µg ml^-1^ DAPI to mid log-grown cells, incubating for 30 min, and washing with PBS before subjecting to imaging. For SINC quantification, cells were grown O/N in complete minimal media, diluted in fresh media to an OD_600_=0.3 and allowed to grow for 4h (∼2 divisions) before imaging. Statistical analyses of the mean values were performed using R statistical language (R Development Core Team, 2008), and different groups are marked with letters at the 0.05 significance level.

### Co-immunoprecipitation

Cell lysates were prepared by resuspending the pellets from 150-200 OD_600_ units in 800 μl lysis buffer (50 mM Tris pH 7.5, 150 mM NaCl, 10% glycerol, 1 mM EDTA pH 8, 0.5% NP-40, 1x complete EDTA-free protease inhibitor cocktail (Roche), 2 mM PMSF, 20 mM N-ethylmaleimide (NEM)). Cells were lysed by bead-beating (Precellys 24, Bertin instruments) with zirconia/silica beads (BioSpec Inc.) and lysates were cleared by centrifugation (800 g, 5 min). Clarified extracts were incubated with antiHA-sepharose (Roche) or GFP-Trap (Chromotek) for 1.5 hours at 4°C, beads were washed four times with lysis buffer and two times with wash buffer (50 mM Tris pH 7.5, 150 mM NaCl, 1 mM EDTA pH 8). Proteins were eluted by boiling with HU loading buffer and analyzed by immunoblotting.

### Protein sequence alignment

Each set of protein sequences was aligned using the MAFFT multiple sequence alignment program (version 6.951b; Katoh and Toh, 2008). The resulting multiple sequence alignments were analyzed for sequence similarities and secondary structures by using the ESPript 3.0 program (Robert and Gouet, 2014).

### Prediction of intrinsically disordered regions

Predictions were performed using the Protein DisOrder prediction System (PrDOS; Ishida and Kinoshita, 2007), with a prediction false positive rate of 2%.

## Acknowledgements

We thank F. Wilfling, C.-W. Lee and members of the Jentsch and Braun labs for fruitful discussions and critical comments on the manuscript; F. Wilfling and C.-W. Lee for strains; and K. Straßer for technical assistance.

Research in the S.J. laboratory was supported by Max Planck Society, Deutsche Forschungsgemeinschaft, Center for Integrated Protein Science Munich (CIPSM), Louis-Jeantet Foundation and a European Research Council (ERC) Advanced Grant. Research in the S.B. laboratory was supported by grants from the German Research Foundation (BR 3511/4-1). S.B. is a member of the Collaborative Research Center 1064 funded by the German Research Foundation and acknowledges infrastructural support.

## Author contributions

MC and SJ conceived the study. MC performed all the experiments, and LMC assisted with the microscopy studies. All authors analyzed the data. MC and SB conceived and wrote the manuscript, and all authors contributed to editing.

## Supplementary information

Fig. S1. Heh1-S localizes to the nuclear envelope and exhibits an epistatic phenotype with NPC genes.

Fig. S2. Genetic interactions between *HEH1* and the ESCRT pathway.

Fig. S3. The MSC domain causes the severe growth phenotype of ESCRT-III mutant cells expressing only Heh1-L.

Fig. S4. Point mutations in the MSC domain of Heh1-L suppress the growth phenotype of ESCRT-III mutants lacking Heh1-S.

Fig. S5. Deletion of *CHM7* rescues the growth defect of ESCRT-III mutants expressing only Heh1-L.

Fig. S6. Heh1-L interacts with Heh1-S.

Fig. S7. Heh1-L and Heh1-S have intrinsically disorder regions mainly at the N-terminal domain.

Table S1. Yeast strains used in this study.

Table S2. Plasmids used in this study.

